# The HMGN Proteins Are Transcriptional Regulatory Factors in Humans

**DOI:** 10.1101/2025.11.27.690916

**Authors:** Grisel Cruz-Becerra, Jonathan Zau, Risa Purow-Ruderman, Ann Dao, Benjamin Delatte, Nathaniel Chapin, George A. Kassavetis, James T. Kadonaga

**Author notes:** Correspondence should be addressed to J.T.K.

## Abstract

The high mobility group N (HMGN) proteins, which were discovered over 50 years ago, are a multigene family of abundant nucleosome-specific binding factors that are present in all vertebrates. Despite their intriguing nucleosome-binding activity, the potential functions of the HMGN proteins in chromatin have not yet been assessed unambiguously due to the presence of several related *HMGN* genes in vertebrates and the lack of *HMGN* null cells. Here, we investigated the genome-wide activities of the human HMGN proteins by generating and analyzing an *HMGN* null cell line and isogenic *HMGN* rescue cell lines. These experiments revealed that the HMGN proteins function in the activation of gene expression at the level of transcription initiation at over a thousand specific sites that are mostly in promoters and enhancers. We additionally observed shared as well as unique functions of HMGN1 and HMGN2, which are likely to be the most abundant and ancient HMGN proteins. These findings thus indicate that the HMGN nucleosome-binding proteins are vertebrate-specific regulatory factors that primarily function in the activation of transcription initiation. Hence, any comprehensive model of vertebrate gene regulation should incorporate the contributions of the HMGN proteins, which are integral components of chromatin in all vertebrates.

## Introduction

The vertebrate-specific HMGN proteins are abundant and ubiquitous non-histone chromosomal factors that bind to two high-affinity sites on nucleosomes in a DNA sequence-independent manner (Goodwin et al. 1973; Mardian et al. 1980; Sandeen et al. 1980; Johns 1982; Ueda et al. 2009; Alegrio-Louro et al. 2025). The HMGN proteins also inhibit ATP-dependent chromatin remodeling (Rattner et al. 2009), probably by competing with the remodeling factors for binding to the nucleosome acidic patch (Alegrio-Louro et al. 2025). Humans contain five different *HMGN* genes (*HMGN1-5*), and the *HMGN1* and *HMGN2* genes have been detected in all vertebrate genomes examined to date (Gonzá-lez-Romero et al. 2015). Remarkably, overexpression of *HMGN1* has been implicated in Down syndrome-associated phenotypes, such as the increased risk of B-cell acute lymphoblastic leukemia and congenital heart defects, in studies with mouse models and patient-derived samples (Mowery et al. 2018; Nanduri et al. 2020; Page et al. 2022; Ranade et al. 2025). In contrast, the effects of the complete loss of the HMGN proteins have not yet been assessed. Prior genetic analyses, which have been carried out nearly entirely in mice or mouse cells, have used partial knockouts that contain at least two unaltered wild-type HMGN genes (Birger et al. 2003; Kugler et al 2013; Deng et al. 2015; Nanduri et al 2020; Page et al. 2022; Abe et al. 2025).

Hence, to elucidate the functions of human HMGN proteins, we deleted all five *HMGN* genes and created an *HMGN* null cell line, with which we generated isogenic *HMGN* rescue cell lines. Then, to gain different perspectives on the activities of the HMGN proteins, we performed genome-wide analyses of gene expression, transcription initiation, and chromatin accessibility. These experiments revealed new and specific insights into the HMGN proteins and demonstrated that they are important components of the transcriptional regulatory system in humans.

## Results

### Generation of *HMGN* null and rescue cells

To examine the functions of the human HMGN proteins, we initially sought to knock out all five of the *HMGN* genes and thus generate an *HMGN* null cell line (Fig. 1). In previous studies, only single or double *HMGN* knockouts had been performed in human cells or mice; hence, the resulting cells or organisms contained multiple functional *HMGN* genes (Birger et al. 2003; Kugler et al. 2013; Deng et al. 2015; Page et al. 2022; Abe et al. 2025). Here, we generated an *HMGN* null cell line by sequentially knocking out the five *HMGN* genes by using CRISPR-Cas9 methodology (Ran et al. 2013) in MCF10A cells (Fig. 1b), which are diploid non-tumorigenic epithelial cells derived from a human mammary gland. The loss of function alleles for all of the *HMGN* genes were determined by DNA sequencing, and the absence of each of the five HMGN proteins in the null cells was assessed by western blot analysis with the parental MCF10A cells (herein referred to as wild-type) as a positive control (Fig. 1c). We also observed that the *HMGN* null cells grew at a rate that is similar to that of the wild-type cells (Extended Data Fig. 1). The viability of the *HMGN* null cells was not unexpected given that non-vertebrates lack HMGN proteins.

**Figure 1.**
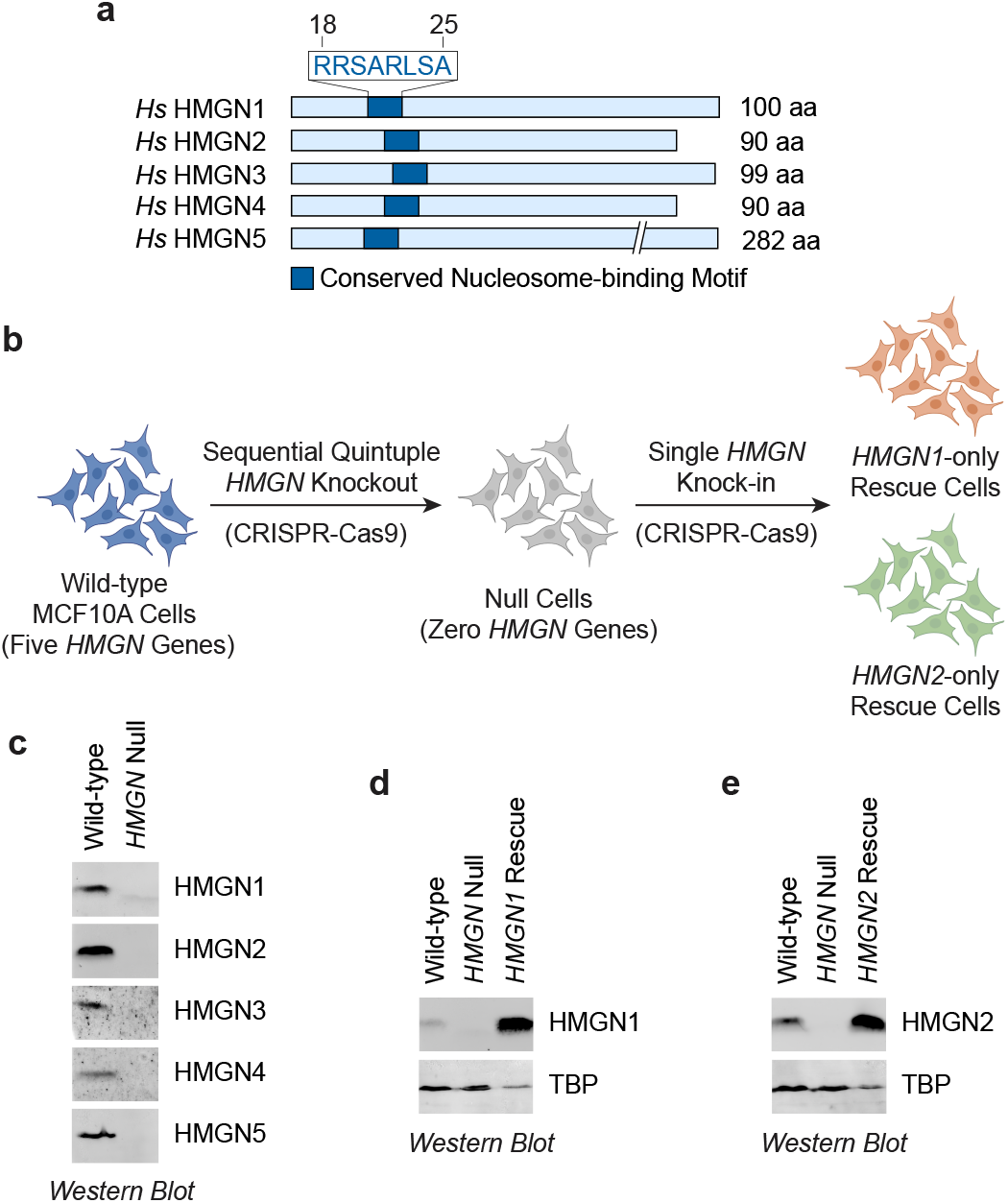
Generation and characterization of human *HMGN* null cells, *HMGN1* rescue cells, and *HMGN2* rescue cells. **a**, Diagram of the five human HMGN nucleosome-binding proteins. The sequence in blue type corresponds to the motif that interacts with the nucleosome acidic patch (Alegrio-Louro et al. 2025) and is present in all five HMGN proteins. **b**, *HMGN* null, *HMGN1* rescue, and *HMGN2* rescue cell lines were generated with human MCF10A cells by using the CRISPR-Cas9 system. Each of the five *HMGN* genes was sequentially knocked out to produce *HMGN* null cells. The coding sequences of HMGN1 and HMGN2 were then separately introduced into the *HMGN* null cells to create rescue lines that express only HMGN1 or HMGN2. **c**, Western blot analysis of wild-type and *HMGN* null cell extracts with HMGN variant-specific antibodies reveals the loss of all five HMGN proteins in the null cells. **d, e**, Detection of HMGN1 and HMGN2 in the rescue cell lines by western blot analysis. HMGN1 and HMGN2 in the rescue lines are expressed at higher levels than in wild-type cells. The lanes with the *HMGN* rescue cell extracts contain one-half of the amount of total protein used in the lanes containing the wild-type and null cell extracts. TBP (TATA box-binding protein) was used as a loading control. Panel b was partially created in BioRender. CRUZ, G. (2025) https://BioRender.com/kkmbqiu.

Next, to enable the unambiguous analysis of the functions of individual HMGN proteins, we used the *HMGN* null cells to generate isogenic rescue cell lines that contain either HMGN1 or HMGN2 (Fig. 1b), which are the most commonly-occurring and likely most ancient HMGN proteins (González-Romero et al. 2015). In this manner, we could determine the functions of the individual HMGN proteins by comparison of the properties of the rescue cells relative to the null cells. The comparison of the isogenic null vs. rescue cells would also exclude any possible off-target effects that might have occurred during the generation of the null cells. Thus, we separately integrated the coding sequences of HMGN1 and HMGN2 into the *AAVS1* locus by using CRISPR-Cas9-mediated homology-directed repair. These experiments resulted in rescue cell lines that express only HMGN1 or HMGN2 in the *HMGN* null genetic background (Fig. 1b, d-e). Hence, the *HMGN* null and rescue cell lines allow, in conjunction with the parental wild-type cells, the precise analysis of the functions of the HMGN proteins.

### HMGN proteins regulate gene expression

The effect of the loss of HMGN proteins on gene expression has not yet been analyzed by the comparison of wild-type vs. null cells. Previous studies used single or double *HMGN* knockouts that contained at least two intact *HMGN* genes (Birger et al. 2003; Rochman et al. 2011; Kugler et al. 2013; Deng et al. 2015). Here, to gain an understanding of the effects of the HMGN proteins upon gene expression, we first performed parallel RNA-seq analyses of wild-type, *HMGN* null, and *HMGN* rescue cells (Fig. 2, Extended Data Fig. 2). These experiments revealed that the loss of all five HMGN proteins affects the expression of 2,016 genes (≥2-fold change; FDR ≤0.01) relative to that seen in the wild-type (control) cells. Among the affected genes, 61% (1,234) have lower expression and 39% (782) have higher expression in the null cells relative to the wild-type cells (Fig. 2a). We then compared the null cells with the *HMGN1* and *HMGN2* rescue cell lines and thus identified shared and HMGN-variant-specific target genes. We examined, in particular, the 483 rescued genes for which there is an opposite effect upon HMGN restoration relative to HMGN loss. This analysis revealed the activation of 305 genes and repression of 178 genes by HMGN1 and/or HMGN2 (Fig. 2b). Somewhat strikingly, over 10 genes were activated >25-fold by HMGN1 and/or HMGN2 (Fig. 2c, Extended Data Fig. 3). The HMGN-dependent genes are predominantly protein-coding genes (87.5–92.1%) along with long non-coding RNAs (7.5–10%) and other gene types (≤2.5%) (Extended Data Fig. 4). *PRR15L* (encoding proline-rich 15-like protein) and *IL24* (encoding interleukin-24 protein) are representative examples of protein-coding genes that are activated and repressed, respectively, by HMGN proteins (Fig. 2d, e). It is also notable that a substantial fraction of the target genes are shared by HMGN1 and HMGN2 (Fig. 2b). These findings suggest that HMGN1 and HMGN2 have overlapping as well as distinct biological functions.

**Figure 2.**
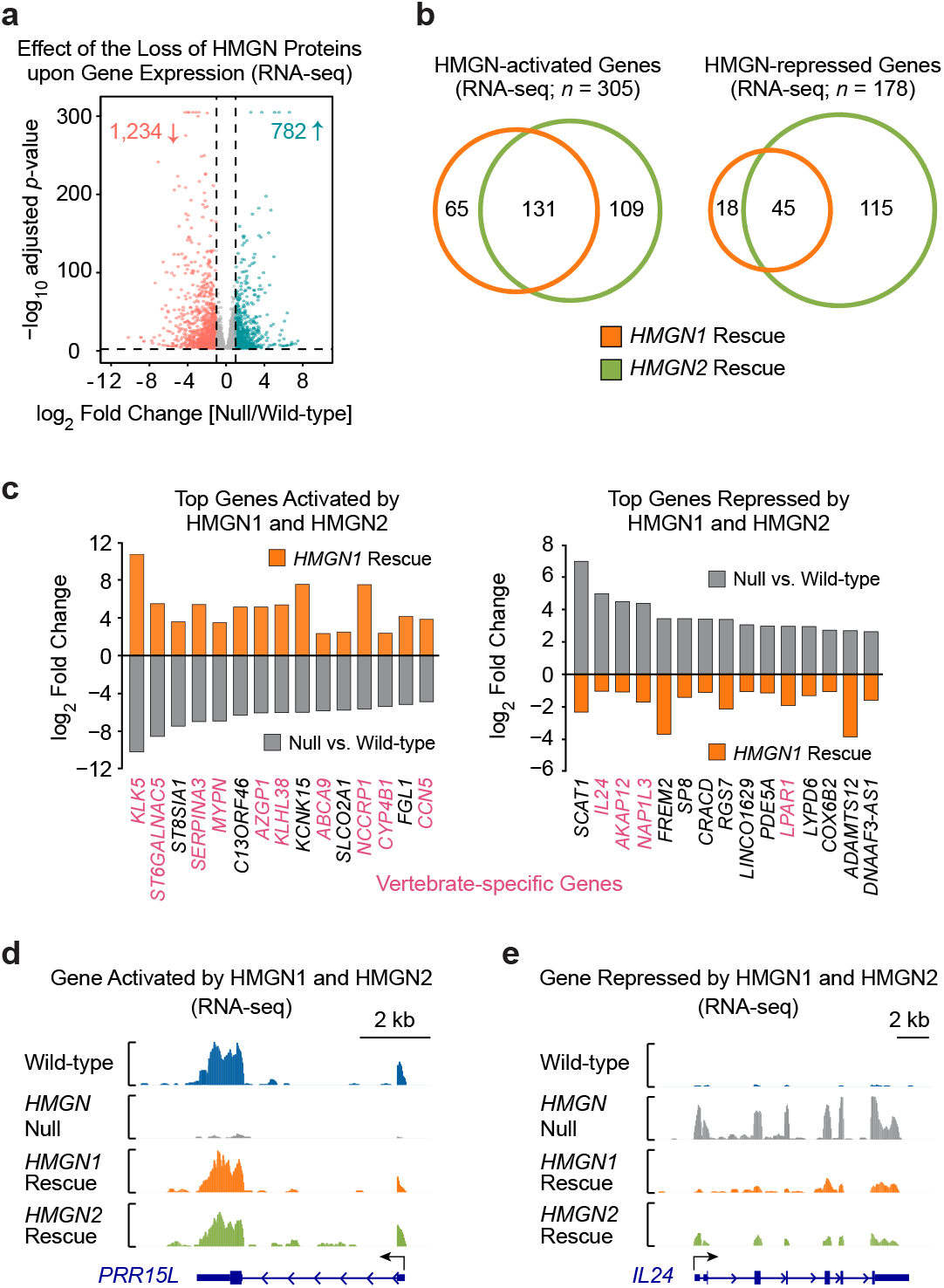
HMGN proteins control the expression of selected genes, including many that are vertebrate-specific. **a**, Volcano plot of differential gene expression (RNA-seq) in wild-type vs. *HMGN* null cells. The loss of all five HMGN proteins results in a ≥2-fold decrease in the expression of 1,234 genes and ≥2-fold increase in the expression of 782 genes. **b**, HMGN1 and HMGN2 activate more genes than they repress. The numbers in the Venn diagrams correspond to genes whose expression is altered by ≥2-fold upon loss of the five HMGN proteins and rescued by ≥2-fold upon re-expression of HMGN1 or HMGN2. **c**, Most of the top genes activated by HMGN1 as well as by HMGN2 are vertebrate-specific genes. Differential expression (log_2_ fold changes in expression for wild-type vs. null and for null vs. rescue) of the top genes activated (left) and repressed (right) by both HMGN1 and HMGN2. Genes that appear to have vertebrate-specific functions are highlighted in pink type. For clarity, only the *HMGN1* rescue data are shown. The corresponding *HMGN2* rescue data are shown in Extended Data Fig. 3. **d, e**, RNA-seq genome browser tracks of representative shared HMGN1- and HMGN2-target genes.

**Figure 3.**
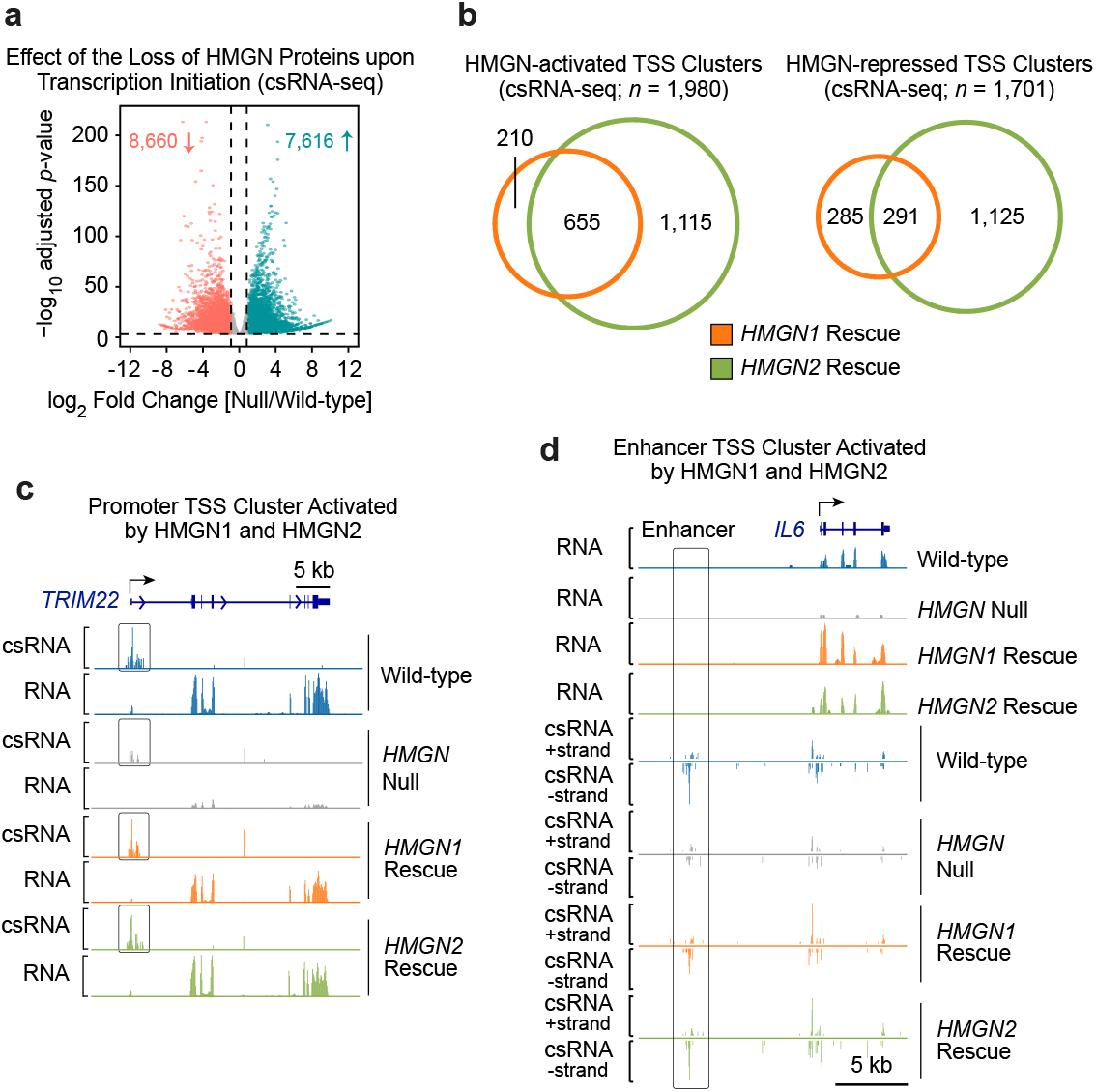
HMGN proteins regulate transcription initiation in human cells. **a**, Volcano plot of changes in active transcription (measured by csRNA-seq; Duttke et al. 2019) in wild-type vs. *HMGN* null cells. The loss of all five HMGN proteins results in a ≥2-fold decrease and a ≥2-fold increase in the activity of 8,660 and 7,616 transcription start site (TSS) clusters, respectively. **b**, HMGN1- and HMGN2-dependent transcription initiation sites. The numbers in the Venn diagrams correspond to TSS clusters whose activity is affected by ≥2-fold upon loss of the five HMGN proteins and rescued by ≥2-fold upon re-expression of HMGN1 or HMGN2. The TSS cluster targets that are shared by HMGN1 and HMGN2 are more frequently activated (655 sites) than repressed (291 sites). **c, d**, Representative promoter (**c**) and enhancer (**d**) TSS clusters that are activated by HMGN1 as well as by HMGN2. The genome browser tracks show RNA-seq (stable RNAs) and csRNA-seq (TSSs of stable and unstable RNAs) data.

**Figure 4.**
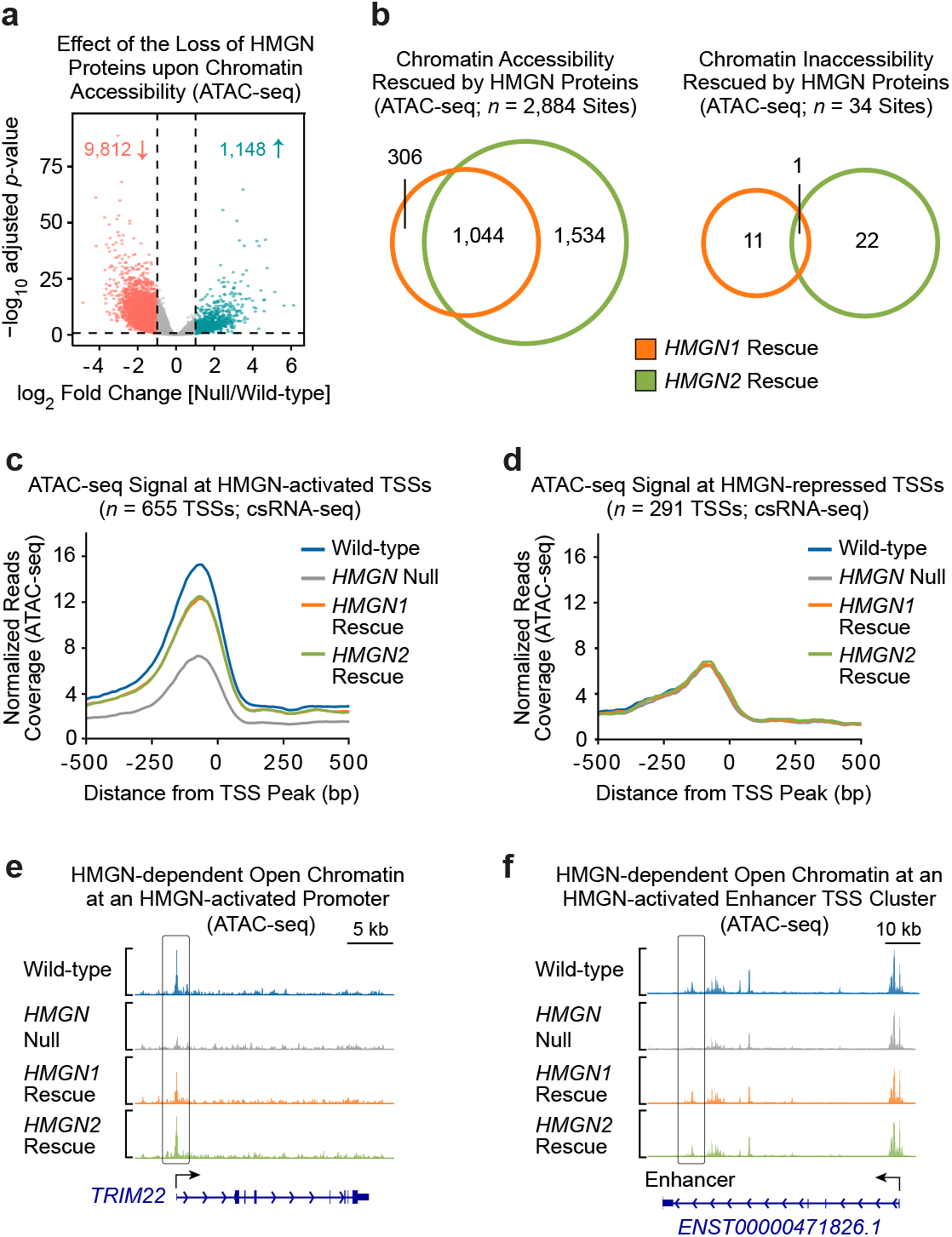
Transcriptional activation but not repression correlates with HMGN-dependent chromatin accessibility. **a**, Volcano plot of differential chromatin accessibility (ATAC-seq) analysis in wild-type vs. *HMGN* null cells. The loss of all five HMGN proteins results in ≥2-fold reduced accessibility at 9,812 genomic sites and ≥2-fold increased accessibility at 1,148 sites. **b**, Re-expression of HMGN1 or HMGN2 in the *HMGN* null cells primarily rescues chromatin opening and has little effect on closed chromatin. The Venn diagrams show the number of genomic sites at which the ATAC signal changes by ≥2-fold upon loss of the five HMGN proteins and is rescued by ≥2-fold upon re-expression of HMGN1 or HMGN2. **c, d**, The presence of HMGN proteins correlates with chromatin accessibility at TSS clusters that are activated by HMGN1 and HMGN2, but not at TSS clusters that are repressed by HMGN1 and HMGN2. The plots in **c** and **d** show normalized ATAC-seq signal (y-axis) from −500 to +500 relative to TSS clusters (at position 0) that are activated (655 TSSs; panel **c**) or repressed (291 TSSs; panel **d**) by HMGN1 and HMGN2. **e, f**, Genome browser views of ATAC-seq signal at a representative promoter (**e**) and enhancer (**f**) activated by HMGN1 and HMGN2.

Intriguingly, many of the most strongly HMGN-activated genes (*i*.*e*., whose expression decreases upon loss of HMGN proteins and increases upon restoration of HMGN proteins) appear to be vertebrate-specific or to have vertebrate-specific functions (Fig. 2c, Extended Data Fig. 3). These genes encode, for instance, factors that are important for lipid metabolism, immune responses, and tissue organization in vertebrates (Bang et al. 2001; Katsube et al. 2009; Jeong et al. 2015; Papenfuss et al. 2015; Dijkstra et al. 2018; Peng et al. 2023). In contrast, we did not observe an enrichment in vertebrate-related genes in the HMGN-repressed targets (Fig. 2c, Extended Data Fig. 3). These findings show that the HMGN proteins regulate the expression of a subset of human genes, and more commonly appear to function in gene activation, particularly of vertebrate-related genes, than in repression.

### HMGN proteins regulate transcription initiation

Because RNA-seq identifies the steady-state reservoir of transcripts, we sought to investigate the effects of HMGN proteins more specifically upon transcription initiation. To this end, we used capped small RNA-seq (csRNA-seq), which detects the transcription start sites (TSSs) of stable and unstable RNAs (Duttke et al. 2019) (Fig. 3, Extended Data Fig. 5). csRNA-seq analysis of wild-type cells vs. *HMGN* null cells showed that the loss of all five HMGN proteins results in a decrease in transcription at 8,660 TSS clusters and an increase in transcription at 7,616 TSS clusters (≥2-fold change; FDR ≤0.01) (Fig. 3a). We then compared the null cells vs. the rescue cells and restricted our analysis to the TSS clusters that exhibited opposite transcriptional effects upon the loss vs. the restoration of HMGN proteins. In this manner, we observed that the re-expression of the HMGN proteins leads to the activation of 1,980 TSS clusters and the repression of 1,701 TSS clusters (Fig. 3b). As seen in the RNA-seq analyses (Fig. 2b), a considerable fraction of the HMGN-regulated TSS clusters are shared by both HMGN1 and HMGN2 (*e*.*g*., ∼33% of activated TSSs) (Fig. 3b). These results further indicate that there are overlapping and unique functions of HMGN1 and HMGN2.

The HMGN-regulated TSS clusters map predominantly to annotated enhancers (enhancer-only; 54.4– 68.2%) and to dually annotated promoter and enhancer regions (22.5–24.2%) (Extended Data Fig. 6) (Fishilevish et al. 2017; Frankish et al. 2023). The HMGN-dependent changes in transcription initiation at promoters and enhancers can be seen at representative individual loci (Fig. 3c, d). The csRNA-seq data thus reveal that there are many HMGN-regulated TSSs and that the HMGN proteins affect gene regulation at the level of transcription initiation. Consistent with these findings, HMGN-regulated TSSs are largely found at promoters and enhancers.

### HMGNs open chromatin at activated TSSs

To investigate whether the HMGN proteins affect chromatin accessibility, we performed parallel ATAC-seq (Buenrostro et al. 2013) experiments with wild-type, *HMGN* null, and *HMGN* rescue cells (Fig. 4, Extended Data Fig. 7). Differential accessibility analysis (≥2-fold change; FDR ≤0.01) between the null and the wild-type cells showed that the loss of all of the HMGN proteins results in a reduction at 9,812 sites and an increase at 1,148 sites (Fig. 4a). We then compared the null vs. the rescue cells and focused on the sites in which HMGN restoration resulted in the opposite effect as that seen upon HMGN loss. In these experiments, we found that the re-expression of HMGN proteins increases chromatin accessibility at 2,884 sites but decreases accessibility at only 34 sites (Fig. 4b). Approximately one-third of the sites at which HMGN proteins increase chromatin accessibility are shared between HMGN1 and HMGN2. In addition, the majority of the sites at which HMGN proteins increase accessibility correlate with annotated enhancers and promoters (Extended Data Fig. 8). Thus, the ATAC-seq analyses indicate that the HMGN proteins increase chromatin accessibility, largely at enhancers and promoters. We next examined whether the effect of the HMGN proteins on chromatin accessibility is associated with their function in transcription initiation. To this end, we quantified the ATAC-seq signals from −500 bp to +500 bp relative to the peaks (position zero) of TSS clusters that are either activated or repressed by HMGN proteins in the rescue of the null cells. These analyses revealed that the HMGN-activated TSS clusters show higher accessibility in the wild-type, *HMGN1* rescue, and *HMGN2* rescue cells than in the *HMGN* null cells (Fig. 4c, e, f, Extended Data Fig. 9a– c). Moreover, the relative amounts of chromatin accessibility at HMGN-activated TSSs in the four cell lines correspond to the relative amounts of HMGN-mediated TSS activation (compare Fig. 4c and Extended Data Fig. 9a–c with Extended Data Fig. 10a). In sharp contrast, chromatin accessibility remains largely unchanged at TSS clusters that are repressed in the *HMGN* null cells (Fig. 4d, Extended Data Fig. 9d–f), and the relative amounts of chromatin accessibility at HMGN-repressed TSSs in the wild-type, null, *HMGN1* rescue, and *HMGN2* rescue cells do not correlate with the relative amounts of HMGN-mediated TSS repression (compare Fig. 4d and Extended Data Fig. 9d–f with Extended Data Fig. 10b). Thus, HMGN-mediated activation of transcription initiation correlates with chromatin opening, but HMGN-mediated repression of transcription initiation does not and is likely to be indirect. These findings collectively lead to a model in which the HMGN proteins function primarily in the activation of transcription initiation at over a thousand specific sites that are mostly in promoters and enhancers.

## Summary and perspectives

The HMGN proteins, which were discovered over 50 years ago (Goodwin et al. 1973), are a family of abundant nucleosome-specific binding proteins that are present in all vertebrates. Here, we investigated the genome-wide functions of the human HMGN proteins. We initially generated isogenic *HMGN* null, *HMGN1* rescue, and *HMGN2* rescue cells, and then analyzed these cells in parallel with the parental wild-type cells. We focused in particular on effects in which HMGN rescue resulted in the opposite effect as HMGN loss. By RNA-seq, we observed that the HMGN proteins regulate a subset of genes, including many with vertebrate-related functions. With csRNA-seq, we found that the HMGN proteins affect transcription initiation at thousands of TSS clusters that are mainly located in promoters and enhancers. Third, in ATAC-seq analyses, we saw that the HMGN proteins increase chromatin accessibility at HMGN-activated TSSs, but not at HMGN-repressed TSSs. These findings suggest that the HMGN proteins function as activators of transcription initiation at promoters and enhancers. In addition, the parallel analyses of the *HMGN* null cells, *HMGN1* rescue cells, and the *HMGN2* rescue cells revealed both shared and unique functions of HMGN1 and HMGN2. Hence, the HMGN proteins contribute a distinct mode of vertebrate-specific transcriptional control alongside regulatory mechanisms such as DNA methylation (Tajima and Suetake 1998; Deaton and Bird 2011; Luo et al. 2018; Klughammer et al. 2023).

This study thus reveals the function of the HMGN proteins in the activation of gene expression at the level of transcription initiation. Therefore, the activities of the HMGN proteins should now be incorporated in the analysis of transcription in vertebrates. Although the transcription process is complex and remains to be clarified, it is nevertheless possible to examine the individual components, including the HMGN proteins, in further detail. Lastly, the genome-wide data that we have generated on HMGN-dependent transcription initiation and chromatin accessibility should serve as a useful resource that would enable researchers to understand how their genes of interest are regulated within a multidimensional framework of the control of gene expression.

## Data availability

The RNA-seq, csRNA-seq, and ATAC-seq data will be made publicly available upon acceptance of this manuscript in a peer-reviewed journal.

## Acknowledgements

We thank Bailey Munro for initial work on HMGN proteins. We are grateful to Active Motif for interactions and collaboration in studies that are separate from but related to in this work. J.T.K. is the Amylin Chair in the Life Sciences. This work was supported by NIH grant R35 GM118060 to J.T.K. G.C.-B. is an UCSD Chancellor’s Fellow and New England Biolabs Fellow. This publication includes data generated at the UC San Diego IGM Genomics Center utilizing an Illumina NovaSeq X Plus that was purchased with funding from a National Institutes of Health SIG grant (#S10 OD026929). This paper was typeset with the bioRxiv word template by @Chrelli: www.github.com/chrelli/bioRxiv-word-template

## Author contributions

G.C.-B. and J.T.K. initially conceived the project and oversaw the overall execution of this work. G.C.-B. performed the genome-wide analyses as well as the computational analyses. R.P.-R., J.Z., G.C.-B., and A.D. generated and/or characterized the knockout cell lines. J. Z. generated the rescue cell lines and performed gene conservation analysis. G.A.K. performed DNA sequence conservation analysis. B.D. and N.C. generated preliminary data. G.C.-B. and J.T.K. prepared the figures and wrote the manuscript.

## Materials and Methods

### Generation and validation of *HMGN* null and rescue cell lines

MCF10A cells, which are non-tumorigenic human epithelial mammary gland cells, were used throughout this study. We purchased the MCF10A cells (which we refer to as the “wild-type”) from the American Type Culture Collection (ATCC). The cells were cultured under standard conditions as reported previously (Cruz-Becerra et al. 2020). *HMGN* null cells were generated by using the CRISPR-Cas9 system described by Ran et al. (2013). Briefly, each of the five *HMGN* genes was knocked out sequentially with a Cas9-expressing plasmid [Addgene plasmid #48138 (a gift from Feng Zhang) or Addgene plasmid #64324 (a gift from Ralf Kuehn)] carrying a crRNA targeting one *HMGN* gene at a time. The crRNAs targeted the first protein-coding exons that contain the conserved HMGN nucleosome-binding motif (Alegrio-Louro et al. 2025). For each knockout, 0.75 million cells per well were seeded in 6-well plates on the day before transfection. The transfection experiments were performed with Lipofectamine 3000 according to the manufacturer’s instructions (Life Technologies). Isogenic clones were derived by single-cell sorting of the transfected cells on an Aria Fusion cell sorter at the Human Embryonic Stem Cell FACS facility, University of California San Diego. All five knockouts were verified by PCR analysis of genomic DNA from isogenic clones by using primers that flank the crRNA target sites. MCF10A cells are diploid with two chromosomal copies of each of the five *HMGN* genes. All ten loss-of-function alleles in the resulting quintuple-knockout cell line were confirmed by Sanger sequencing, and the absence of all five HMGN proteins was confirmed by western blot analysis. To generate rescue cell lines that contain only HMGN1 or HMGN2, we used homology-directed repair-mediated CRISPR-Cas9 integration at the *AAVS1* safe-harbor locus in the *HMGN* null cells. Donor DNA templates contained the HMGN1 or HMGN2 coding sequence and a promoterless neomycin/Geneticin (G418) resistance cassette, and they were co-transfected with a Cas9-expressing plasmid bearing an *AAVS1*-targeting crRNA sequence. Transfection and isogenic clone isolation were as described in the knockout workflow described above, except that the edited cell pools were first selected with Geneticin before single-cell sorting. The expression of HMGN1 or HMGN2 in the rescue cell lines was confirmed by western blot analysis with variant-specific antibodies (Cell Signaling Technology).

### Cell growth analysis

Growth curves for wild-type (MCF10A) and *HMGN* null cell lines were determined by using the crystal violet assay as described by Feoktistova et al. (2016). Briefly, 4,000 cells per well were seeded in 96-well plates for both cell lines. Separate plates were prepared for each time point (*i*.*e*., 1, 2, 3, 5, 6, and 7 days) at which crystal violet staining was quantified. The absorbance of crystal violet at 570 nm was measured by using the Tecan Spark® Microplate Reader.

### RNA-seq experiments and data analysis

RNA-seq experiments were performed with three biological replicates for each cell line. Total RNA was prepared by using TRIzol Reagent according to the manufacturer’s instructions (Invitrogen). The RNA samples (6 µg per reaction) were incubated with DNase I (4 units) for 20 minutes at 37 °C in 1x DNase I reaction buffer (100 µL final reaction volume) (New England Biolabs; NEB). The reactions were stopped by the addition of 1 µL of 0.5 M EDTA (Na^+^), pH 8.0, to give a final concentration of 5 mM EDTA. The RNA was subsequently purified by using the Monarch Spin RNA Cleanup Kit according to the protocol recommended by the manufacturer (NEB). The RNA quality was assessed by using an Agilent 4200 Tapestation, and samples with an RNA Integrity Number (RIN) ≥9.8 were used to generate sequencing libraries with the Illumina Stranded mRNA Prep (Illumina). The samples were processed according to the manufacturer’s instructions, and the resulting libraries were sequenced with paired-end 150-bp reads to a depth of approximately 50 million reads per sample on an Illumina NovaSeq X Plus sequencer at the IGM Genomics Center, University of California, San Diego. Adapter sequence trimming and low-quality read filtering were performed with fastp (v1.0.1; Chen et al. 2018). The resulting reads were aligned to the GRCh38 (hg38) reference genome by using STAR (v2.7.10a; Dobin et al. 2013). Reads mapping to exons were quantified with FeatureCounts (Subread v2.0.3; Liao et al. 2014) by using the GENCODE release 48 annotation. Differential expression between wild-type and *HMGN* null cells as well as between *HMGN* null and rescue cells was assessed with DESeq2 (Love et al. 2014). Genes with log_2_ fold change ≥1 (absolute) and an adjusted *p*-value <0.01 were considered significant.

### csRNA-seq experiments and data analysis

The csRNA-seq experiments were performed with a minimum of two biological replicates for each cell line. Total RNA was prepared by using TRIzol Reagent according to the manufacturer’s instructions (Invitrogen). csRNA and input libraries were prepared by using total RNA as described in Duttke el al. (2019) and sequenced with paired-end 150-bp reads to a depth of approximately 25 million reads per sample on an Illumina NovaSeq X Plus sequencer at the IGM Genomics Center, University of California, San Diego. The sequencing data were analyzed according to the Christopher Benner laboratory (University of California, San Diego) csRNA-seq analysis pipeline (http://homer.ucsd.edu/homer/ngs/csRNAseq/) with HOMER (Heinz et al. 2010; Duttke et al. 2019) with the following modifications. To call valid TSSs, findcsRNATSS.pl in HOMER was run with data from matched input, csRNA, and RNA libraries for individual replicates by using a minimum read count (-ntagThreshold) of 10 (measured in reads per 10 million). Peaks were removed from the data if a RepeatMasker element overlapped the 50-bp window centered on the TSS summit (≥1-bp overlap). Only the clusters that passed the filter and were reproducible in ≥2 replicates for the same condition were retained for differential expression analysis with DESEq2 (Love et al. 2014). TSS clusters with log_2_ fold change ≥1 (absolute) and an adjusted *p*-value <0.01 were considered significant. GENCODE release 48 (Frankish et al 2023; GENCODE Project) and GeneHancer data (Fishilevish et al. 2017) were used to annotate the TSSs genomic features.

### ATAC-seq experiments and data analysis

ATAC-seq experiments were performed with two biological replicates per cell line. ATAC libraries were prepared from 80,000 freshly collected cells per replicate by using the Active Motif ATAC-seq Kit according to the manufacturer’s protocol (Active Motif). The nucleosomal laddering was assessed on an Agilent 4200 TapeStation, and the resulting libraries were sequenced as paired-end 50-bp reads to a depth of approximately 60 million reads per sample on an Illumina NovaSeq X Plus at the IGM Genomics Center, University of California, San Diego. Adapters were trimmed and low-quality reads were filtered with fastp (v1.0.1; Chen et al. 2018). The resulting reads were aligned to the human hg38 reference genome by using Bowtie2 (v2.4.5; Langmead and Salzberg 2012). Mitochondial alignments and reads with low-quality alignment scores were removed prior to assesing library complexity, and PCR duplicates were then removed with Picard (v2.18.29; Broad Institute). To correct the Tn5 transposase offset, alignments were shifted +4 bp on the forward strand and −5 bp on the reverse strand by using deepTools AlignmentSieve (v3.5.1; Ramirez et al. 2016). MACS2 (v2.2.9.1; Zhang et al. 2008) was used to call peaks for individual replicates with a false discovery rate <0.05 and then regions overlapping the ENCODE blacklist were removed (annotation ENCSR636HFF; Amemiya et al. 2019; ENCODE Project Consortium). The filtered peaks that were reproducible in the two replicates for the same condition were retained for differential accessibility analysis with DiffBind (Stark and Brown 2011), comparing wild-type vs. *HMGN* null and *HMGN* null vs. rescue data. Sites with log_2_ fold change ≥1 (absolute) and an adjusted *p*-value <0.01 were considered significant.

## EXTENDED DATA

**Extended Data Fig. 1.**
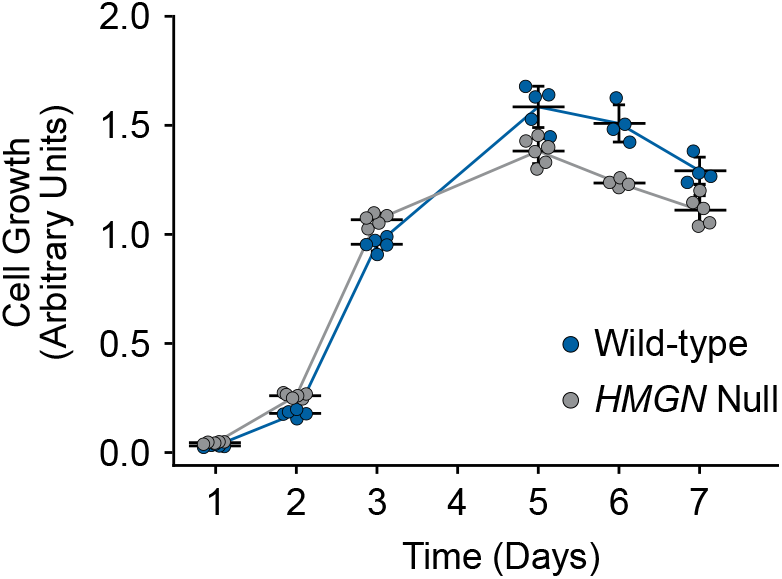
*HMGN* null cells grow at a rate that is similar to that of the wild-type. The scatter plot shows the absorbance at 570 nm by crystal violet staining of *HMGN* null and wild-type cell lines over 7 days, with measurements every 24 hours. All wells were seeded at equal initial cell density, and each time point included ≥4 technical replicates. The two cell lines grow at comparable rates and reach a plateau after 5 days.

**Extended Data Fig. 2.**
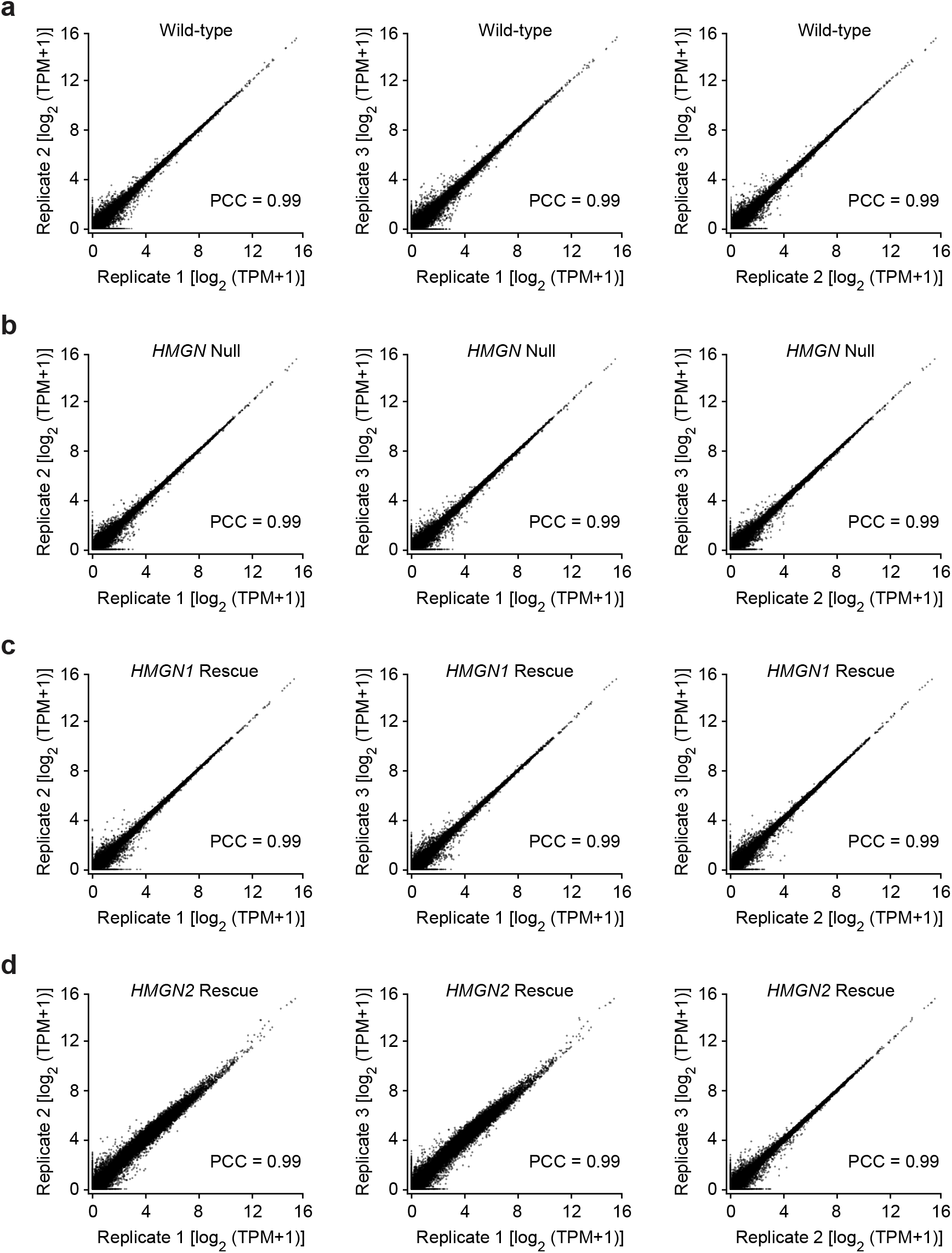
The RNA-seq data are highly reproducible in biological replicates for each condition. The scatter plots show pairwise comparisons of gene expression [log2(TP-M+1)] for three biological replicates with wild-type (**a**), *HMGN* null (**b**), *HMGN1* rescue (**c**), and *HMGN2* rescue (**d**) cell lines. TPM, transcripts per million; PCC, Pearson correlation coefficient.

**Extended Data Fig. 3.**
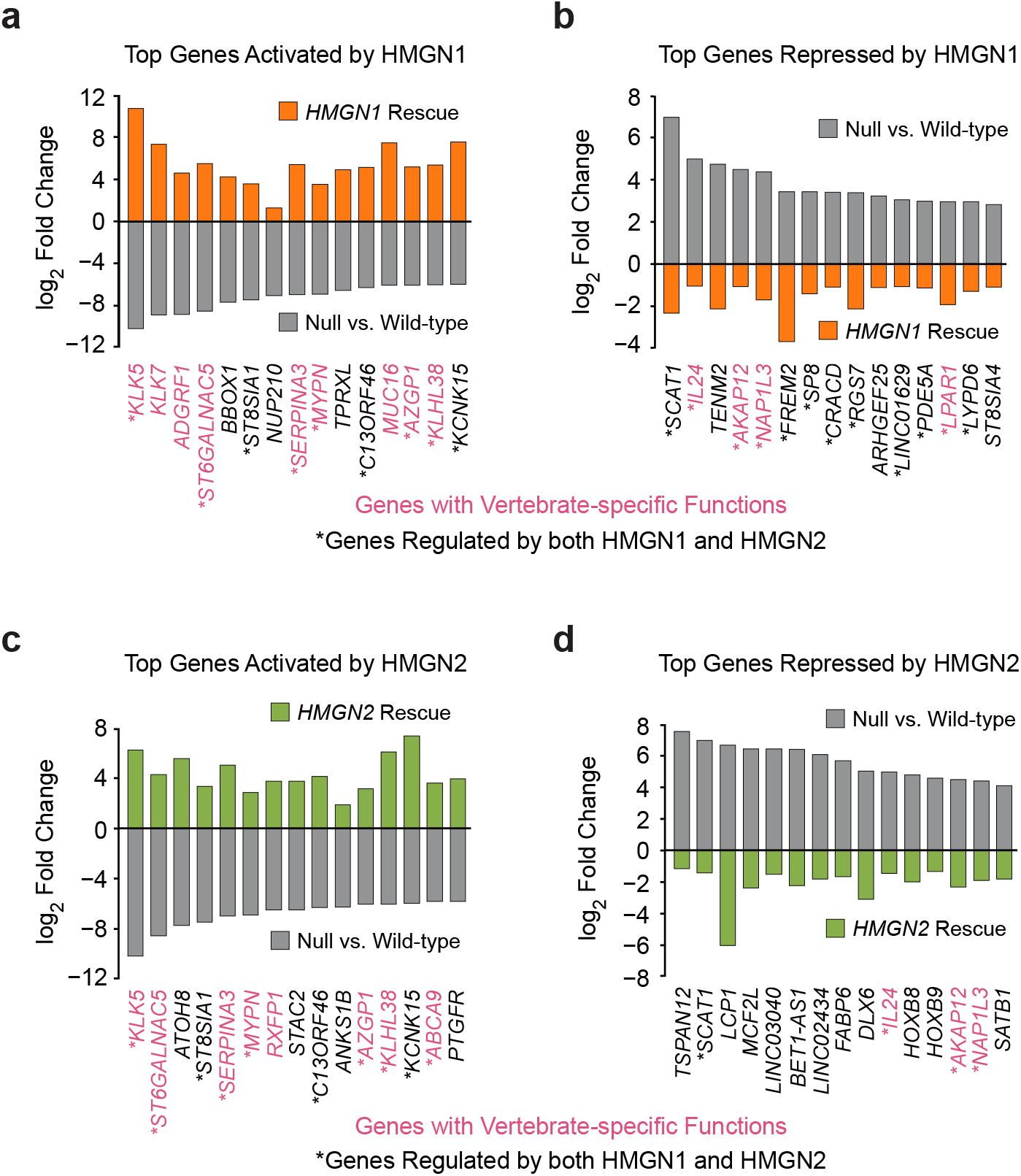
Top target genes activated or repressed by HMGN1 or HMGN2. **a-d**, Each panel shows the top 15 genes whose expression (as assessed by RNA-seq) is altered by ≥ 2-fold upon loss of the five HMGN proteins and rescued by ≥2-fold upon re-expression of HMGN1 (**a, b**) or HMGN2 (**c, d**). The graphs show the log_2_ fold changes in expression for wild-type vs. null (gray bars) and for null vs. rescue (orange bars for *HMGN1* rescue; green bars for *HMGN2* rescue). The asterisks indicate shared target genes between HMGN1 and HMGN2. Genes that appear to have vertebrate-specific functions are highlighted in pink type.

**Extended Data Fig. 4.**
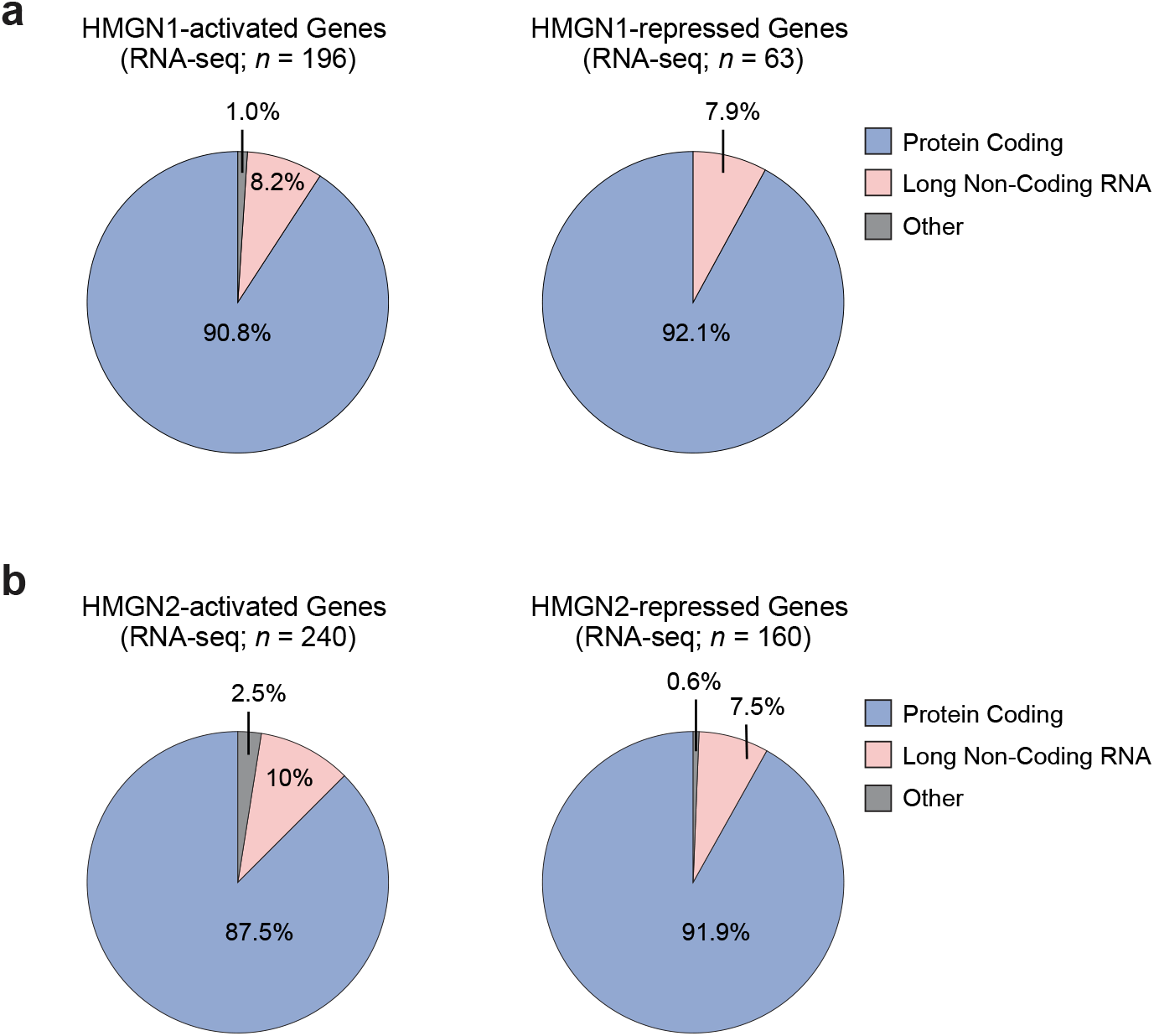
Activation and repression by HMGN1 and HMGN2 predominantly affect protein-coding genes. **a, b**, The pie charts show the most frequent biotype classes (GENCODE annotation release 48) of the genes that are activated or repressed (as assessed by RNA-seq) by HMGN1 (**a**) or HMGN2 (**b**). Protein-coding is the predominant class (≥87.5%) across the four gene sets. Among the HMGN1-activated genes (*n* = 196), 90.8% are protein-coding, 8.2% are long non-coding RNA, and 1.0% belong to other classes. Within the HMGN1-repressed genes (*n* = 63), 92.1% and 7.9% are protein-coding and long non-coding RNA, respectively. Among the HMGN2-activated genes (*n* = 240), 87.5% are protein-coding, 10.0% are long non-coding RNA, and 2.5% belong to other classes. Within the HMGN2-repressed genes (*n* = 160), 91.9% are protein-coding, 7.5% are long non-coding RNA, and 0.6% (one gene) belongs to a different class. The “other” class includes putative protein-coding genes and processed pseudogenes.

**Extended Data Fig. 5.**
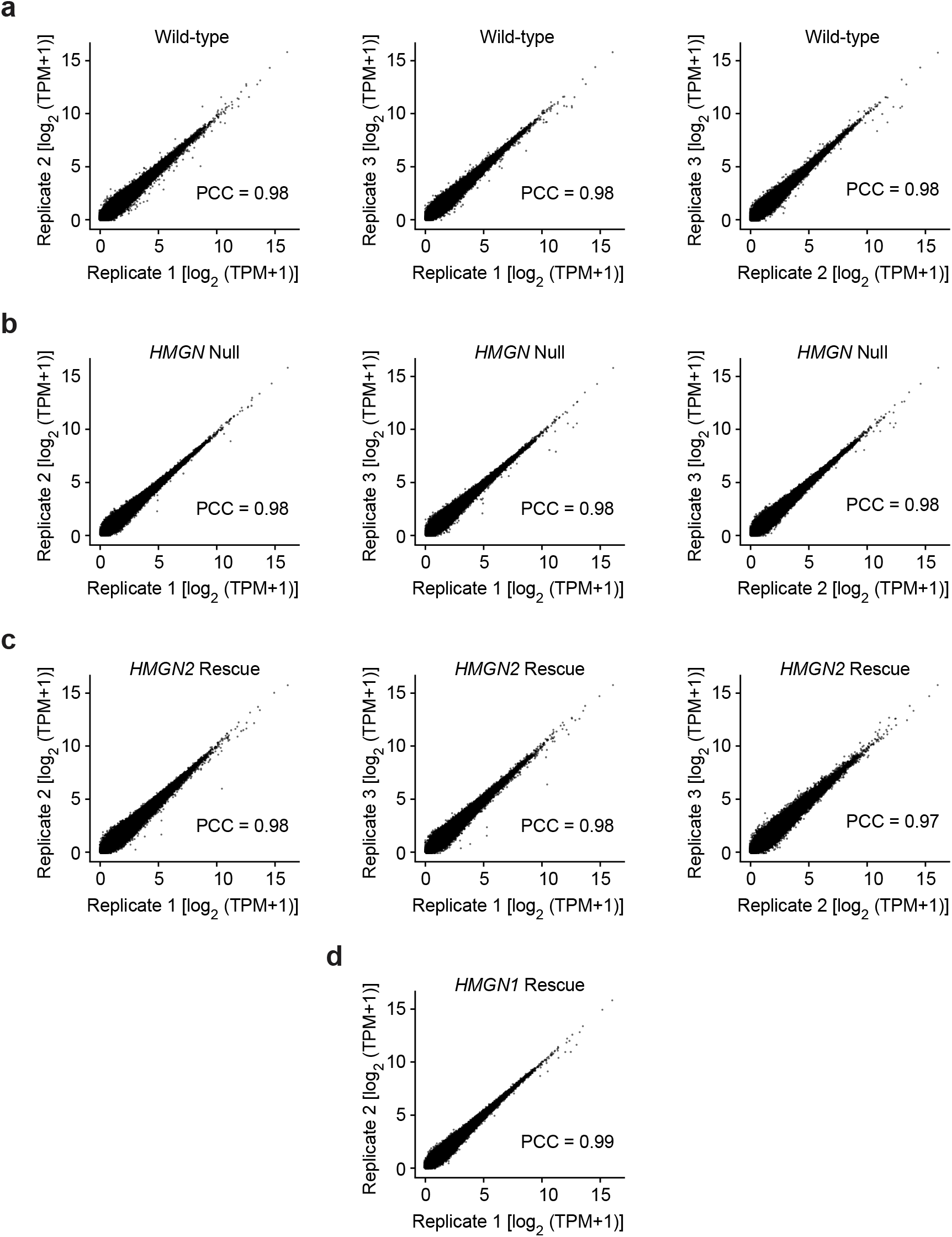
The csRNA-seq data are highly reproducible in biological replicates for each condition. The scatter plots show pairwise comparisons of the TSS signals [log2(TP-M+1)] for two or three biological replicates with wild-type (**a**), *HMGN* null (**b**), *HMGN2* rescue (**c**), and *HMGN1* rescue (**d**) cell lines. TPM, transcripts per million; PCC, Pearson correlation coefficient.

**Extended Data Fig. 6.**
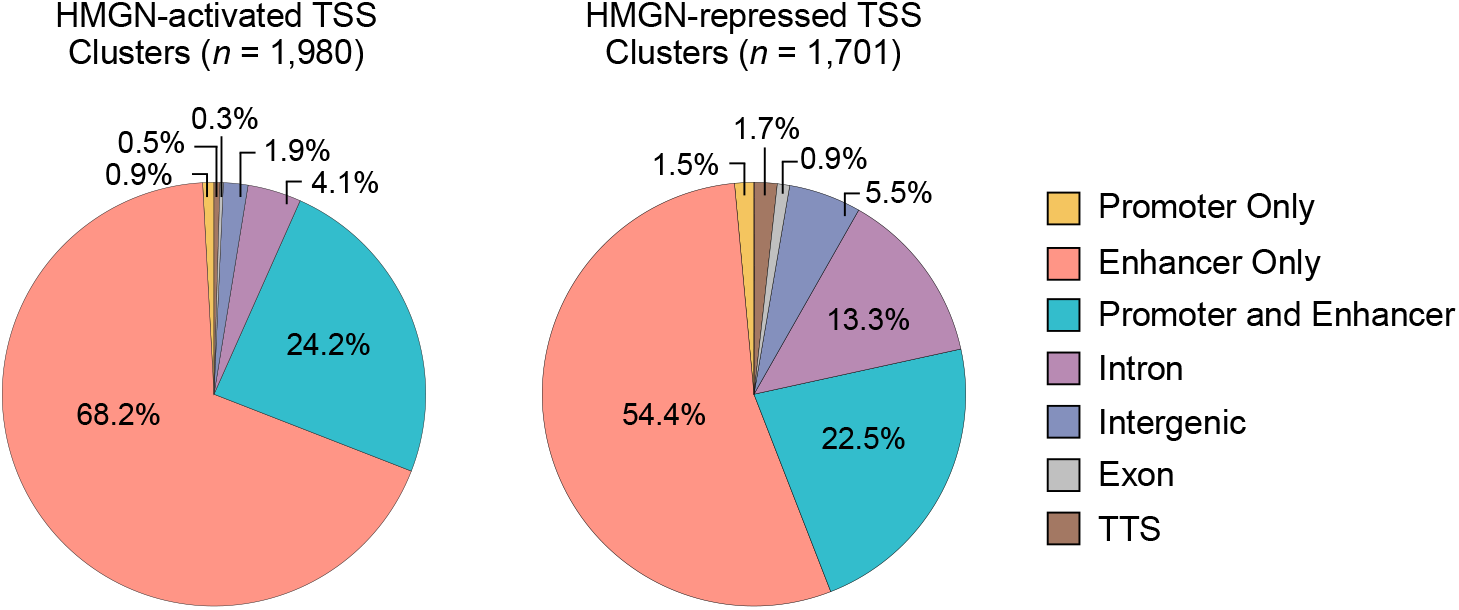
HMGN1 and HMGN2 regulate transcription initiation at gene promoters and enhancers. The TSS cluster targets that are activated (1,980 clusters) or repressed (1,701 clusters) by HMGN1 and/or HMGN2 predominantly map to annotated enhancers and to regions that are classified as both enhancers and promoters (Fishilevish et al. 2017; Frankish et al. 2023).

**Extended Data Fig. 7.**
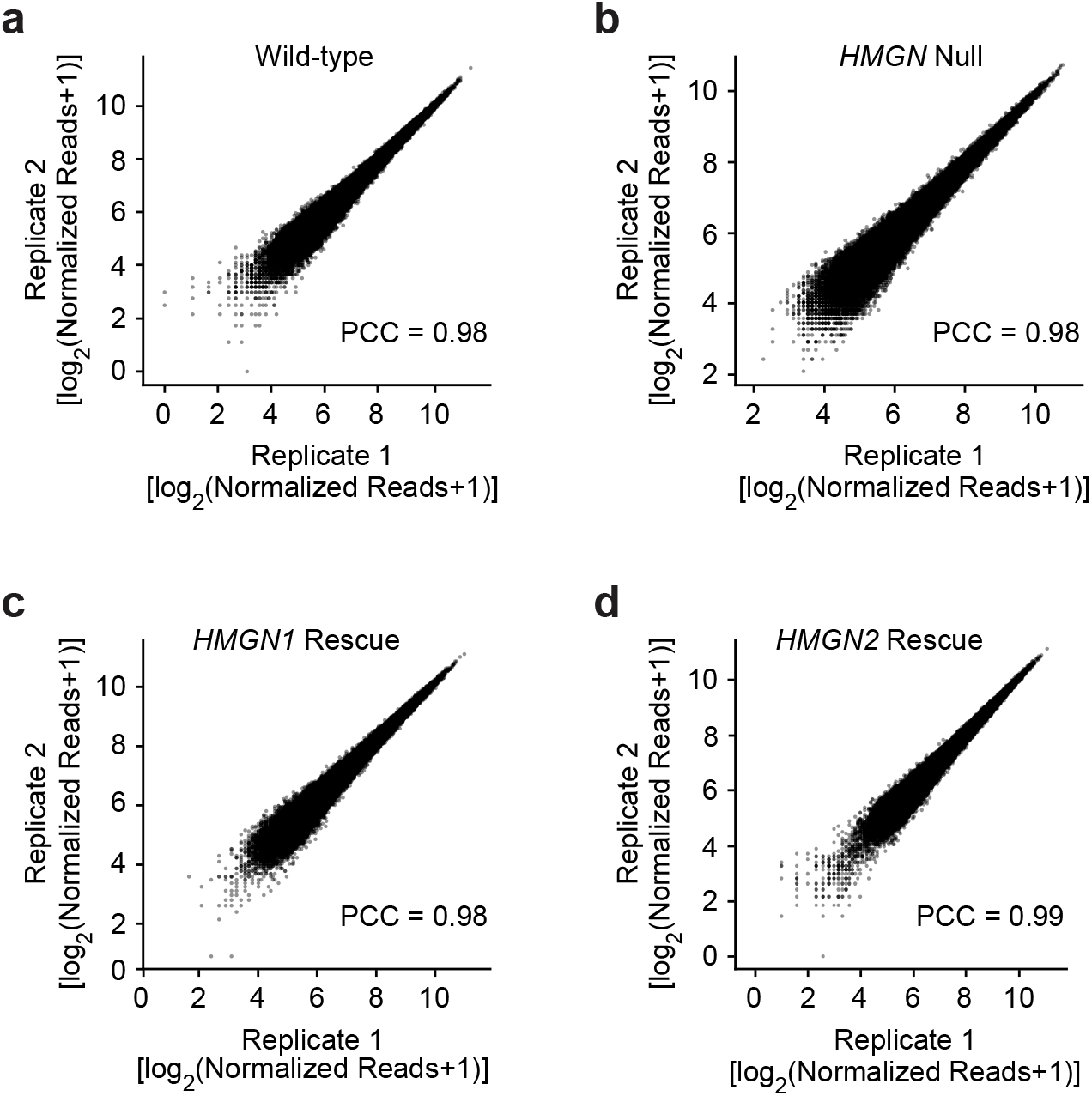
The ATAC-seq data are highly reproducible in biological replicates for each condition. The scatter plots show pairwise comparisons of normalized read coverage [log2(normalized read counts+1)] for two biological replicates with wild-type (**a**), *HMGN* null (**b**), *HMGN1* rescue (**c**), and *HMGN2* rescue (**d**) cell lines. TPM, transcripts per million; PCC, Pearson correlation coefficient.

**Extended Data Fig. 8.**
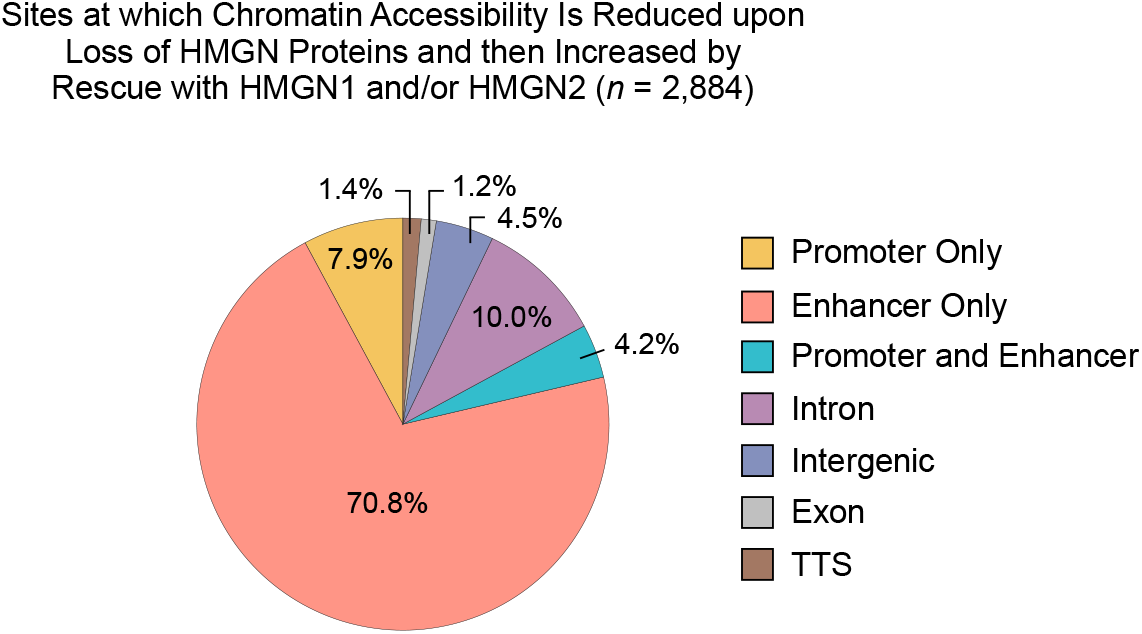
HMGN proteins are associated with open chromatin, predominantly at enhancers. Re-expression of HMGN1 and/or HMGN2 in the *HMGN* null cells increases accessibility at 2,884 sites that had lost accessibility in the null cells relative to wild-type cells. The majority of these sites map to annotated enhancers (Fishilevish et al. 2017; Frankish et al. 2023).

**Extended Data Fig. 9.**
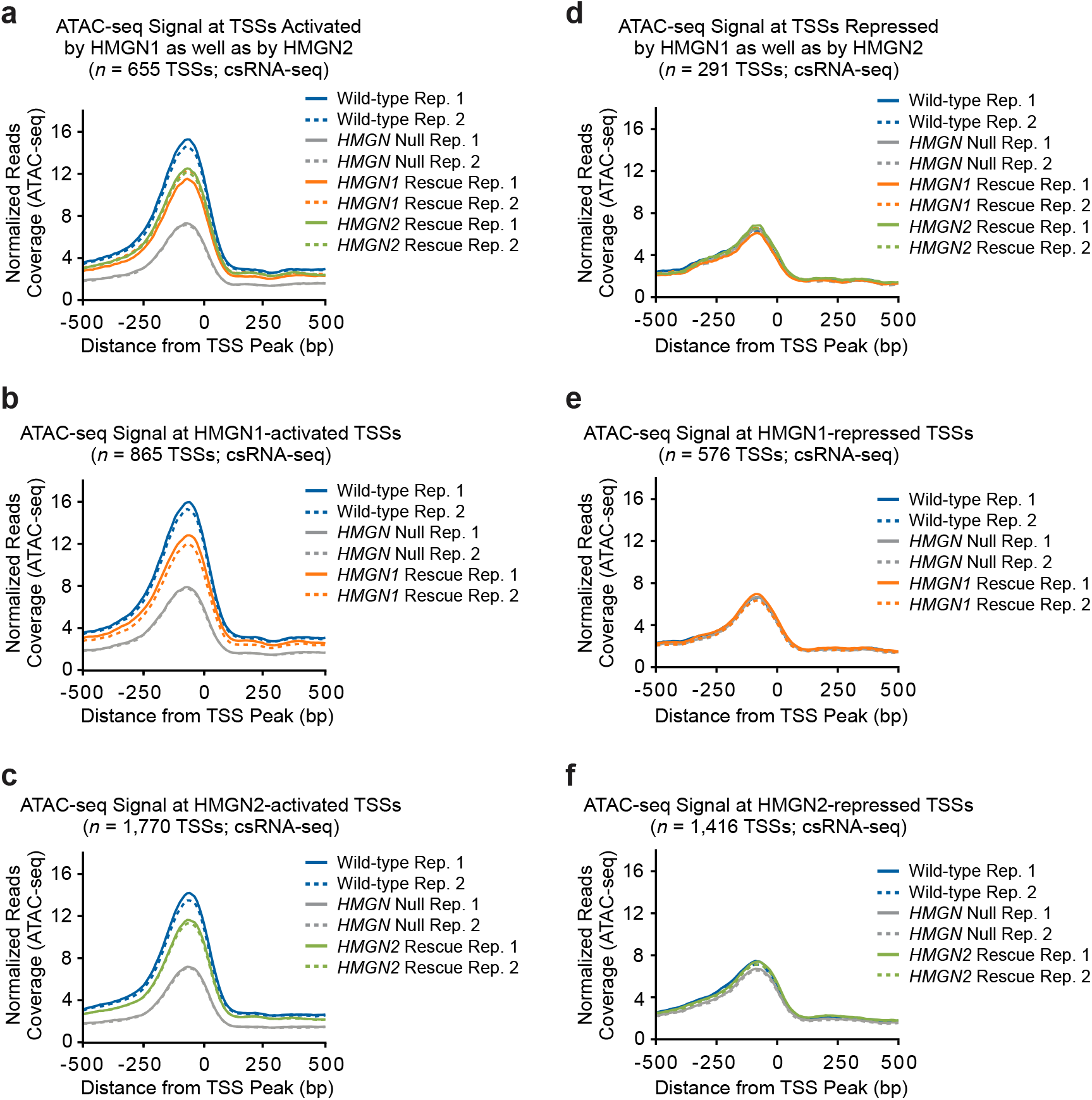
The presence of HMGN proteins correlates with chromatin accessibility at HMGN-activated TSSs, but not at HMGN-repressed TSSs. Normalized ATAC-seq signal (y-axis) from −500 to +500 relative to TSS clusters (at position 0) that are activated or repressed by HMGN proteins. The plots show data for two biological replicates (Rep. 1 and Rep. 2) for each condition. **a–c**, The presence of HMGN proteins correlates with ATAC-seq signal at HMGN1- and HMGN2-activated genes. **d–f**, The presence of HMGN proteins does not correlate with ATAC-seq signal at HMGN1- and HMGN2-repressed genes.

**Extended Data Fig. 10.**
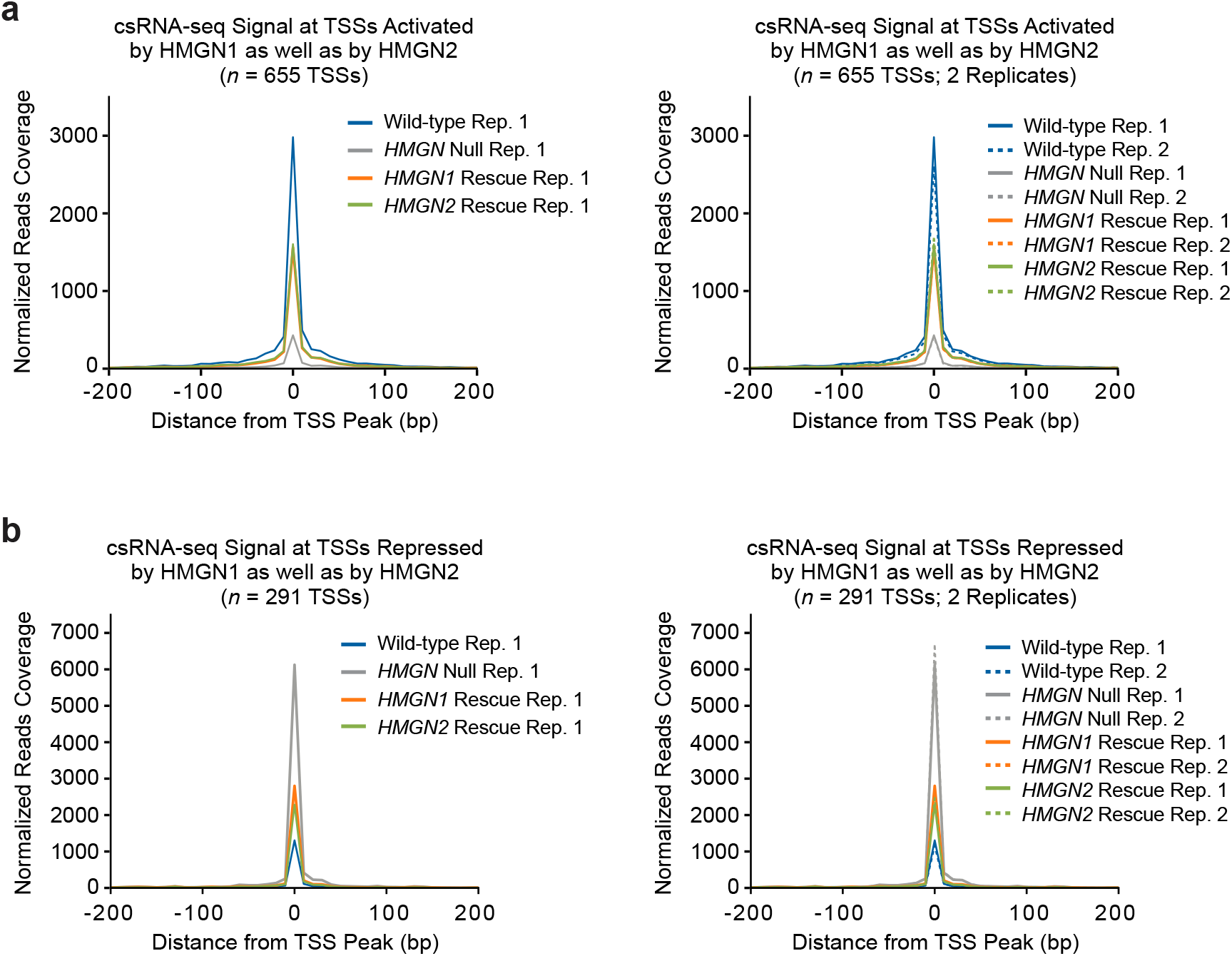
The presence of HMGN proteins correlates with csRNA signals of HMGN-activated TSS clusters and inversely correlates with csRNA signals of HMGN-repressed TSS clusters. The plots show the normalized csRNA-seq signal (y-axis) from −200 to +200 relative to TSS clusters (at position 0) that are activated (*n* = 655) or repressed (*n* = 291) by HMGN1 and HMGN2. For clarity, the panels on the left contain data for one biological replicate for each condition. The panels on the right show that the data are highly reproducible between biological replicates. **a**, The csRNA signal at HMGN1- and HMGN2-activated TSSs correlates with the presence of HMGN proteins. **b**, The csRNA signal at HMGN1- and HMGN2-repressed TSSs inversely correlates with the presence of HMGN proteins.

## References

1. Abe, T. et al. Retinal pigment epithelium specific metabolic phenotypes are regulated by high-mobility group protein N1. Invest ophthalmol & vis sci. 66, 70 (2025).

2. Alegrio-Louro, J., Cruz-Becerra, G., Kassavetis, G. A., Kadonaga, J. T. & Leschziner, A. E. Structural basis of nucleosome recognition by the conserved Dsup and HMGN nucleosome-binding motif. Genes Dev. 39, 1155–1161 (2025).

3. Amemiya, H. M., Kundaje, A. & Boyle, A. P. The ENCODE Blacklist: Iden-tification of problematic regions of the genome. Sci. Rep. 9, 9354 (2019).

4. Bang, M. L. et al. Myopalladin, a novel 145-kilodalton sarcomeric protein with multiple roles in Z-disc and I-band protein assemblies. J. Cell Biol. 153, 413–427 (2001).

5. Birger, Y. et al. Chromosomal protein HMGN1 enhances the rate of DNA repair in chromatin. EMBO J. 22, 1665–1675 (2003).

6. Buenrostro, J. D., Giresi, P. G., Zaba, L. C., Chang, H. Y. & Greenleaf, W. J. Transposition of native chromatin for fast and sensitive epigenomic profiling of open chromatin, DNA-binding proteins and nucleosome position. Nat. Methods 10, 1213–1218 (2013).

7. Chen, S., Zhou, Y., Chen, Y. & Gu, J. fastp: an ultra-fast all-in-one FASTQ preprocessor. Bioinformatics 34, i884–i890 (2018).

8. Chu, V. T. et al. Increasing the efficiency of homology-directed repair for CRISPR-Cas9-induced precise gene editing in mammalian cells. Nat. Biotechnol. 33, 543–548 (2015).

9. Cruz-Becerra, G. & Kadonaga, J. T. Enhancement of homology-directed repair with chromatin donor templates in cells. eLife 9, e55780 (2020).

10. Deaton, A. M. & Bird, A. CpG islands and the regulation of transcription. Genes Dev. 25, 1010–1022 (2011).

11. Deng, T. et al. Functional compensation among HMGN variants modulates the DNase I hypersensitive sites at enhancers. Genome Res. 25, 1295– 1308 (2015).

12. Dijkstra, J. M., Yamaguchi, T. & Grimholt, U. Conservation of sequence motifs suggests that the nonclassical MHC class I lineages CD1/PROCR and UT were established before the emergence of tetrapod species. Immunogenetics 70, 459–476 (2018).

13. Dobin, A. et al. STAR: ultrafast universal RNA-seq aligner. Bioinformatics 29, 15–21 (2013).

14. Duttke, S. H., Chang, M. W., Heinz, S. & Benner, C. Identification and dynamic quantification of regulatory elements using total RNA. Genome Res. 29, 1836–1846 (2019).

15. Feoktistova, M., Geserick, P. & Leverkus, M. Crystal violet assay for determining viability of cultured cells. Cold Spring Harb. Protoc. 2016, pdb.prot087379 (2016).

16. Fishilevich, S. et al. GeneHancer: genome-wide integration of enhancers and target genes in GeneCards. Database (Oxford) 2017, bax028 (2017).

17. Frankish, A. et al. GENCODE: reference annotation for the human and mouse genomes in 2023. Nucleic Acids Res. 51, D942–D949 (2023).

18. González-Romero, R., Eirín-López, J. M. & Ausió, J. Evolution of high mobility group nucleosome-binding proteins and its implications for vertebrate chromatin specialization. Mol. Biol. Evol. 32, 121–131 (2015).

19. Goodwin, G. H., Sanders, C. & Johns, E. W. A new group of chromatin-associated proteins with a high content of acidic and basic amino acids. Eur. J. Biochem. 38, 14–19 (1973).

20. Heinz, S. et al. Simple combinations of lineage-determining transcription factors prime cis-regulatory elements required for macrophage and B cell identities. Mol. Cell 38, 576–589 (2010).

21. Jeong, C. B. et al. Marine medaka ATP-binding cassette (ABC) superfamily and new insight into teleost Abch nomenclature. Sci. Rep. 5, 15409 (2015).

22. Johns, E. W. (ed.) The HMG Chromosomal Proteins Academic Press, London (1982).

23. Kasinsky, H. E., Lewis, J. D., Dacks, J. B. & Ausió, J. Origin of H1 linker histones. FASEB J. 15, 34–42 (2001).

24. Katsube, K., Sakamoto, K., Tamamura, Y. & Yamaguchi, A. Role of CCN, a vertebrate specific gene family, in development. Dev. Growth Differ. 51, 55–67 (2009).

25. Klughammer, J. et al. Comparative analysis of genome-scale, base-resolution DNA methylation profiles across 580 animal species. Nat Commun. 14, 232 (2023).

26. Kugler, J. E. et al. High mobility group N proteins modulate the fidelity of the cellular transcriptional profile in a tissueand variant-specific manner. J. Biol. Chem. 288, 16690–16703 (2013).

27. Langmead, B. & Salzberg, S. L. Fast gapped-read alignment with Bowtie 2. Nat. Methods 9, 357–359 (2012).

28. Liao, Y., Smyth, G. K. & Shi, W. featureCounts: an efficient general-purpose program for assigning sequence reads to genomic features. Bioinformatics 30, 923–930 (2014).

29. Love, M. I., Huber, W. & Anders, S. Moderated estimation of fold change and dispersion for RNA-seq data with DESeq2. Genome Biol. 15, 550 (2014).

30. Luo, C., Hajkova, P. & Ecker, J. R. Dynamic DNA methylation: In the right place at the right time. Science 361, 1336–1340 (2018).

31. Mardian, J. K., Paton, A. E., Bunick, G. J. & Olins, D. E. Nucleosome cores have two specific binding sites for nonhistone chromosomal proteins HMG 14 and HMG 17. Science 209, 1534–1536 (1980).

32. Mowery, C. T. et al. Trisomy of a Down Syndrome critical region globally amplifies transcription via HMGN1 overexpression. Cell Rep. 25, 1898–1911.e5 (2018).

33. Nanduri, R., Furusawa, T. & Bustin, M. Biological functions of HMGN chromosomal proteins. Int J Mol Sci. 21, 449 (2020).

34. Page, E. C. et al. HMGN1 plays a significant role in CRLF2 driven Down Syndrome leukemia and provides a potential therapeutic target in this high-risk cohort. Oncogene 41, 797–808 (2022).

35. Papenfuss, A. T. et al. Marsupials and monotremes possess a novel family of MHC class I genes that is lost from the eutherian lineage. BMC Genomics 16, 535 (2015).

36. Pavlopoulou, A., Pampalakis, G., Michalopoulos, I. & Sotiropoulou, G. Evolutionary history of tissue kallikreins. PLoS One 5, e13781 (2010).

37. Peng, F. et al. Genomic and transcriptional profiles of Kelch-like (klhl) gene family in polyploid Carassius complex. Int. J. Mol. Sci. 24, 8367 (2023).

38. Picard toolkit. Broad Institute, GitHub repository. Broad Institute Available at: http://broadinstitute.github.io/picard/

39. Ramírez, F. et al. deepTools2: a next generation web server for deep-sequencing data analysis. Nucleic Acids Res. 44, W160–W165 (2016).

40. Ran, F. A. et al. Genome engineering using the CRISPR-Cas9 system. Nat. Protoc. 8, 2281–2308 (2013).

41. Ranade, S.S. et al. Myocardial reprogramming by HMGN1 underlies heart defects in trisomy 21. Nature (2025).

42. Rattner, B. P., Yusufzai, T. & Kadonaga, J. T. HMGN proteins act in opposition to ATP-dependent chromatin remodeling factors to restrict nucleosome mobility. Mol. Cell 34, 620–626 (2009).

43. Rochman, M. et al. Effects of HMGN variants on the cellular transcription profile. Nucleic Acids Res. 39, 4076–4087 (2011).

44. Sandeen, G., Wood, W. I. & Felsenfeld, G. The interaction of high mobility proteins HMG14 and 17 with nucleosomes. Nucleic Acids Res 8, 3757– 3778 (1980).

45. Stark, R. & Brown, G. DiffBind: differential binding analysis of ChIP-Seq peak data. Bioconductor (2011).

46. Tajima, S. & Suetake, I. Regulation and function of DNA methylation in vertebrates. J Biochem. 123, 993–999 (1998).

47. Ueda, T., Catez, F., Gerlitz, G. & Bustin, M. Delineation of the protein module that anchors HMGN proteins to nucleosomes in the chromatin of living cells. Mol Cell Biol. 28, 2872–2883 (2008).

48. Zhang, Y. et al. Model-based analysis of ChIP-Seq (MACS). Genome Biol. 9, R137 (2008).

